## Supplementary Data for "A computational toolbox to investigate the metabolic potential and resource allocation in fission yeast"

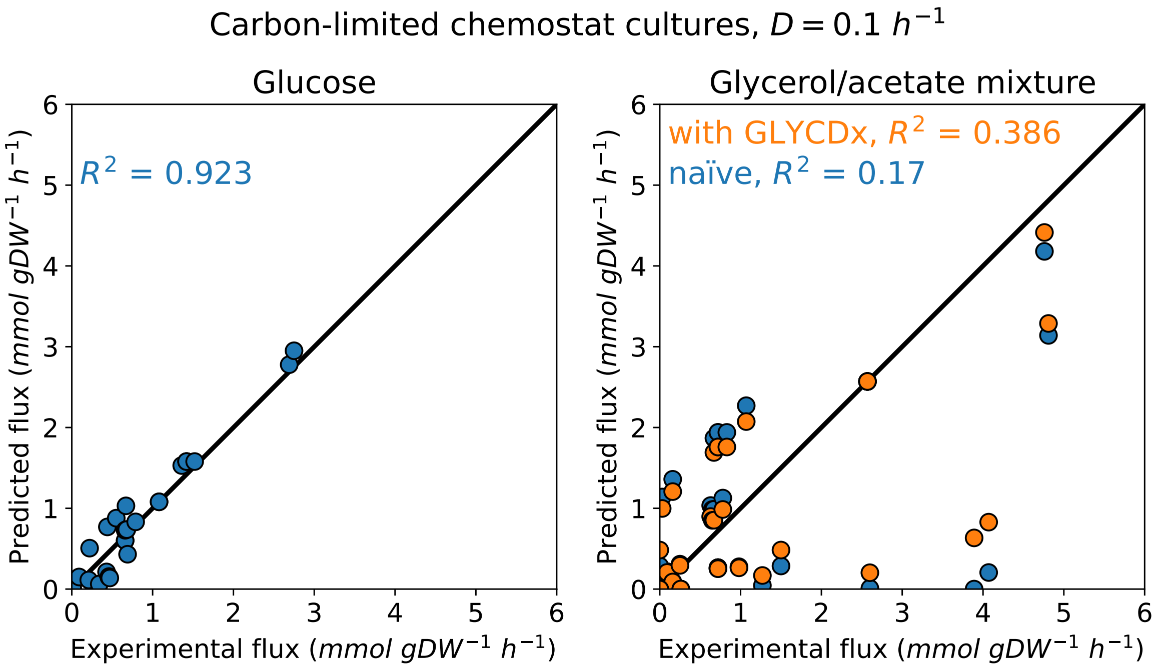

**Supplementary Figure** 1: Flux distributions in carbon-limited chemostat cultures with glucose (left) and glycerol/acetate mixture (right) as carbon sources. *pomGEM* predictions (absolute values of fluxes) plotted on the Y-axis, compared to the experimentally determined values (X-axis). $^{13}C$ labeling data taken from [Klein et al., 2013].

In the case of growth on a glycerol/acetate mixture, a potential model curation step is shown: in the *pomGEM* model, enzyme glycerol dehydrogenase (EC 1.1.1.156) uses $NADP^+$ as the co-factor (BiGG reaction identifier *GLYCDy*). Addition of a $NAD^+$-dependent reaction (*GLYCDx*) resulted in improved agreement with experimental data.

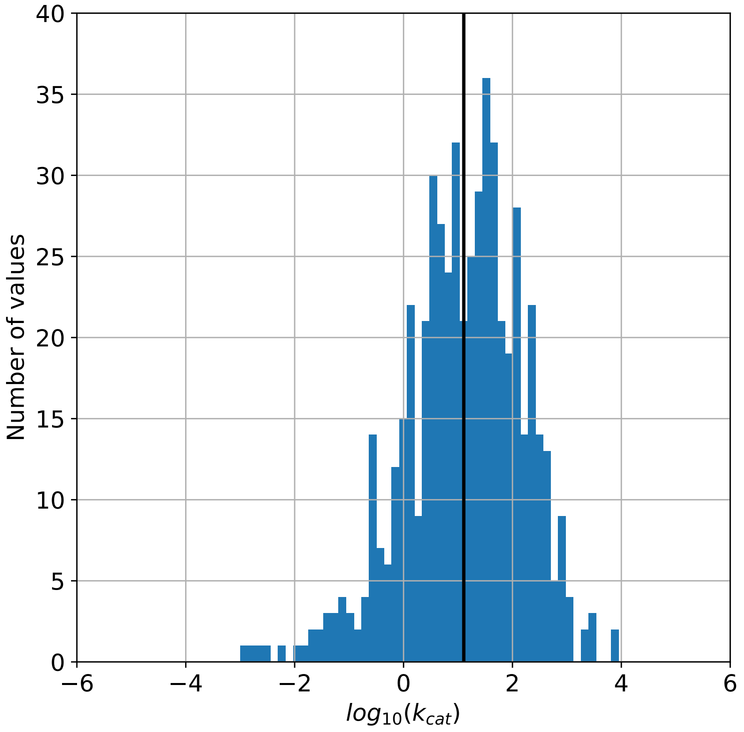

**Supplementary Figure** 2: Distribution of turnover values $k_{cat}$, collected for the *pcPombe* model. Thick black line represents the median $k_{cat}$ value.

### pcPombe model documentation

Pranas Grigaitis *et al.*

April 11, 2022

#### Overview

This document summarizes the construction of the proteome-constrained model of *Schizosaccharomyces pombe*, the *pcPombe*. The *pcPombe* model is a computable knowledge-base, storing of 3 types of descriptions in terms of reaction stoichiometries and constraints: (1) metabolic reactions (the *pomGEM* model part), (2) macromolecule turnover and usage, and (3) proteome capacity constraints for different compartments.

### 1 Modifications of the metabolic model

#### 1.1 Amino acid turnover in mitochondria

The central expansion of the conventional genome-scale model *pomGEM* is introduction of fine-grained descriptions of protein synthesis, folding and degradation. In yeasts, ca. 20 proteins are encoded in the mitochondrial genome and are translated by the mitochondrial ribosome. Therefore, the descriptions of amino acid turnover in mitochondria should be complete as well: we need to define (1) amino acid transport in and out of mitochondria and (2) charging of amino acids onto transport RNAs (tRNAs). In the initial reconstruction of the *pomGEM* model, however, this was not the case. Therefore we reviewed these processes in the *pomGEM* model and updated it accordingly.

###### Amino acid transport

Mitochondrial transporters for many amino acids were not present in the *pomGEM* model, or were defined as amino acid uniport (however, the correct mode of amino acid transport to mitochondria is symport with protons [Wipf et al., 2002]). We have removed all existing amino acid transport reactions, and created new reactions with the following generic form: AA[c]+$H^+$[c] $\leftrightarrow$ AA[m]+$H^+$[m]. Different amino acid permeases were assigned to these transport reactions: for glycine, *SPAC823.10c* and *SPAC4G9.20c* were assigned as transport proteins, and in the case of glutamate and serine, associated transporters were *SPAC4G9.20c* and *SPAC17G6.15c*, respectively. For the rest of amino acids, the low-specificity amino acid permease *SPBC460.01c* was set to catalyze the transport (all annotations collected from PomBase [Lock et al., 2019]).

###### Amino acid-tRNA synthases

For some amino acids (Ala, Arg, Cys, Gly, Lys, Gln, Pro, and Ser), reactions of tRNA charging were missing. If needed, we first created free and charged tRNAs species to the model. Next, we added amino acid-tRNA synthase reactions with the following generic form: AA[m] + tRNA(AA)[m] + ATP[m] $\rightarrow$ AA-tRNA(AA)[m] + AMP[m] + $PP_i$[m].

#### 1.2 Biomass equation in *pcPombe*

In the *pcPombe* model, we use the original biomass equation of the *pomGEM* model (*R_Biomass*), and modify the stoichiometric coefficients of different biomass components to reflect the growth rate $\mu$-dependent changes in biomass composition.

##### 1.2.1 Modifications to the original biomass equation

###### Removal of reactions

The biomass equation had, among its products, a species called "Biomass" (*M_Biomass*), which were removed from the system in a dedicated reaction "Growth" (*R_Growth*) (*M_Biomass* $\rightarrow \emptyset$). Both the species *M_Biomass* and reaction *R_Growth* were removed from the model. Non-growth-associated maintenance (NGAM) reaction *R_Maintenance* was also removed, as ATP maintenance costs in *pcPombe* are a part of the flux through *R_Biomass*.

###### Removal of protein pool metabolite

In the *pomGEM* model, the protein demand for the biomass is defined as "uncharging" of charged amino acid-tRNAs, retrieving the free tRNAs. However, the *pcPombe* model explicitly accounts for proteome turnover, thus the biomass equation itself should not include dilution of proteins by growth. Therefore we removed the reagent *M_Protein* from the reaction *R_Biomass*.

##### 1.2.2 Growth rate-dependent biomass composition

The abundance of some biomass components in *S. pombe* cells is known to be growth rate-dependent. For these biomass components, we recompute the coefficients in the biomass equation based on the growth rate. A summary of the growth rate-relations of different biomass components can be found in Table 1.

Table 1: Growth rate-dependent biomass composition. Data collected from [de Jong-Gubbels et al., 1996, Agar and Bailey, 1981, de Queiroz et al., 1993]

| Component | Reaction | Species | Relation |
|---|---|---|---|
| **Lipids** | *R_Phospholipid* | *M_Phospholipid* | 1 |
| **Carbohydrates** | *R_Carbohydrates* | *M_Carbohydrates* | 0.872 - $m_{RNA}$ - $m_{Prot}$ |
| **DNA** | *R_DNA* | *M_DNA* | $m_{DNA} = 0.0397$ |
| **RNA** | *R_RNA* | *M_RNA* | $m_{RNA} = 0.0812 \times \mu + 0.08222$ |
| **Cofactors** | *R_COF* | *M_COF* | 0.011037527 |
| **Protein** | | | $m_{Prot} = 0.43$ |
| **ATP**<br>GAM part<br>NGAM part | <br>*R_Biomass*<br>*R_Biomass* | | <br>$6.0\ mmol\ gDW^{-1} \times \mu$<br>$0.19\ mmol\ gDW^{-1}\ h^{-1}$ |

#### 1.3 Mitochondria-specific maintenance reaction

In order to more precisely account for the energetic costs, specific to maintaining mitochondria (e.g. costs of protein transport, turnover of nucleic acids etc.), we added an additional reaction to the model, $v_{MitoMaintenance}$. In the current implementation, the substrate of this reaction is mitochondrial ATP (Table 2). Unlike the biomass equation, the stoichiometric coefficients in this equation are scaled to gram of mitochondrial protein, rather than gram of dry cell weight. For this, we coupled the flux through this reaction to the mitochondrial protein content, using a coupling constraint (see Section 3.1.2). As the abundances of individual mitochondrial proteins are optimization variables, so, subsequently, is the flux through this reaction.

Table 2: Components of mitochondrial maintenance reaction as a function of mitochondrial proteome mass $m_{MitoProtein}$.

| Component | Stoich. coefficient |
|---|---|
| **ATP** (hydrolysis) | -6.0 |

#### 1.4 Minor fixes

##### Adding missing metabolic reactions for growth on minimal medium

The growth on the *pomGEM* model was mainly tested on the rich YES medium. As a result, some reactions, essential for biomass formation on a minimal EMM2 medium, were missing, thus we added the missing metabolic reactions to the model in order to fill the remaining gaps (Table 3).

Table 3: Metabolic reactions, added to the *pomGEM* model.

| Reaction ID | Name | GPR |
|---|---|---|
| *R_SRR* | Serine racemase | *SPCC320.14* |
| *R_PHPYR_D* | Phenylpyruvate decarboxylase | *SPAC3G9.11c* |
| *R_PPAmt* | Inorganic pyrophosphatase, mitochondrial | *SPAC3A12.02* |
| *R_ADKmt* | Adenylate kinase, mitochondrial | *SPAC4G9.03* |
| *R_NADH2_u6mm* | NADH dehydrogenase, mitochondrial | *SPBC947.15c* |

##### Reversibility of glycerol:$H^+$ symporter

In the *Yeast8* model (which was used as a template for *pomGEM*), the mode of glycerol transport through plasma membrane depends on the direction: the uptake of glycerol to the cell was modeled as glycerol:$H^+$ symport (*r*_1171, *pomGEM*: glycerol uniporter *R_GLYCt*) [Ferreira et al., 2005], and its export was set as uniport "via channel" (*r*_1172, *pomGEM*: *R_GLYCt2*). Both of these reactions are set as irreversible, suggesting that only one mode is active in one direction. To resolve this issue, we set the glycerol transport reaction $R\_GLYCt$ to be a reversible glycerol:$H^+$ symporter reaction, and removed the reaction $R\_GLYCt2$, as well as the resulting orphan gene *SPAC24H6.01c* from the model.

##### Stoichiometry of monosaccharide import

In *S. cerevisiae*, most of the monosaccharides are known to be imported via facilitated diffusion. In *S. pombe*, transport of glucose is glucose:$H^+$ symport with the stoichiometry of 1:0.4 [Höfer and Nassar, 1987]. Thus, in the *pomGEM* and *pcPombe* models, reactions of glucose transport (*GLCt*) were set to be proton symport by adding 0.4 $H^+[e]$ and 0.4 $H^+[c]$ as reaction substrate and product, respectively.

##### Proton export coupling

In most of the GEMs, mitochondrial ATP synthase reaction is defined as transport of cytosolic protons into mitochondria, irregardless of the "origin" of these protons. Thus, additional influx of protons by glucose transport might be used to generate extra ATP in the mitochondria (strictly speaking, only protons pumped out by respiratory chain complexes should be used for ATP synthesis). This issue is especially important at low growth, where a small increment in ATP production would result in a substantial increase of the predicted growth rate. To adequately account for the costs of keeping balance of protons due to the glucose:$H^+$ import, we set the equal amount of protons to be exported. For this, we couple the fluxes through $H^+$-ATPase (*PMA1*) and plasma membrane glucose transporters (*PMc*, with the glucose:$H^+$ stoichiometry of 1:0.4 [Höfer and Nassar, 1987]):

$$\sum_{i \in PMA1} v_i - 0.4 \sum_{j \in PMc} v_j = 0 \tag{1}$$

##### Transport of organic acids

Plasma membrane transporters of 4 organic acids (acetate, malate, succinate, and fumarate) were defined as uniporters, however, they have been reported as acid:$H^+$ symporters (acetate) or antiporters (malate, succinate, and fumasrate) in *S. cerevisiae* ([Casal et al., 1996, Cassio et al., 1987, Sousa et al., 1992]). However, the main transporter of dicarboxylic acids in *S. pombe* (*mae1*) is defined as proton symport protein in the PomBase. Therefore, all the corresponding transporters ($R\_ACtr$, $R\_MALt$, $R\_SUCCt$, $R\_FUMtr$) were set as 1:1 acid:$H^+$ symporters. The GPR annotations for the permease of dicarboxylic acids were missing, thus gene *SPAPB8E5.03* (*mae1*) was added to the model and assigned to these reactions.

#### 1.5 Splitting of metabolic reactions

##### Splitting `OR` gene-protein-reaction associations

If a gene-protein-reaction (GPR) association did not have an `OR` relationship, then we just used the current GPR association in order to update the reaction name (added the required protein names as a suffix to the reaction name, reaction name *R1* becomes *R1___GPR*). If the reaction GPR had an `OR` relationship, the reaction was deep-copied and new reactions with individual GPR strings were created. E.g.: GPR of a reaction *R1* `(G1 OR (G2 and G3) OR G4)` was split into reactions *R1___G1*, *R1___G2*, *R1___G3*, and *R1___G4*.

##### Splitting reversible reactions

In the *pc*-models, reactions which have proteins associated to them must be unidirectional (otherwise, a reverse reaction would be either impossible and/or would produce metabolic enzymes). Here we split such reactions into two forward and reverse reactions. E.g.: a reversible reaction *R1* was split into forward and reverse reactions *R1_fwd* and *R1_rev*, respectively.

### 2 Description of proteome turnover and reaction coupling

We created different model species for each protein entity of the UniProt reference proteome of *S. pombe* (*UP000002485*) [UniProt Consortium, 2020] in the model, for a sample species with the UniProt accession ID *P00000*:

- Unfolded protein species (*P00000_uf_c*)
- Folded protein species (*P00000_c*)
- "Used" protein species (*P00000_used_c*)

The target compartment in the example is cytoplasm (species suffix "*_c*"), and for proteins, translated in mitochondria, or transported to compartments other than cytoplasm, we created corresponding model species as well. The turnover of proteins in the model (conversions from one species to another) can be described through several classes of reactions, which we will describe in the next sections.

For implementing enzyme coupling constraints, we use "protein complex" species. These species represent the enzymatic complexes (could consist of a single protein as well) at their stoichiometric ratios (if known, otherwise assumed to be 1). A complex formation reaction of $cplx_i$, $v_{cplx\ formation,\ i}$ takes a general form, where complex is formed from individual folded protein species $j$:

$$v_{cplx\ formation,\ i} : \sum_{j \in cplx_i} n_j\ j \rightarrow cplx_i \tag{2}$$

The flux through reactions which need $cplx_i$ for catalysis are coupled with the flux through $v_{cplx\ formation,\ i}$. To maintain the steady-state assumption, two options of consuming the "formed" complex are provided: degradation of the complex into individual *used* proteins $j\_used$ (Eq. 3), and dilution by growth (Eq. 4). In steady-state, $v_{cplx\ formation,\ i} = v_{cplx\ degradation,\ i} + v_{cplx\ dilution,\ i}$. The latter two processes, respectively, are represented as follows:

$$v_{cplx\ degradation,\ i} : cplx_i \rightarrow \sum_{j \in cplx_i} n_j\ j\_used \tag{3}$$

$$v_{cplx\ dilution,\ i} : cplx_i \rightarrow \emptyset \tag{4}$$

The ratio between the flux through complex degradation and dilution-by-growth reactions is growth rate $\mu$-dependent, and is defined in the following section.

#### 2.1 General form of coupling constraints

The use of macromolecular complexes (alike metabolic enzymes and protein turnover machinery [e.g., ribosomes]) is coupled with the fluxes through reactions they catalyze. We describe this relationship between enzyme demand and metabolic flux as follows:

$$v = [e] \times k_{cat} \times f(\mathbf{x}, T, \dots) \tag{5}$$

Where $f$ is a saturation function (with a range $[0; 1]$), which is dependent on different parameters, such as metabolite concentrations, temperature etc. In the *pc*-models, we assume $f = 1$, i.e. enzymes working at their maximal speed. This way, we predict the *minimal* enzyme demand, needed to sustain the metabolic flux. For each enzyme complex, we then couple the flux through complex formation reaction to the sum of fluxes, which that complex catalyzes, scaled with the $k_{cat}$ values:

$$\frac{v_{cplx\ formation,\ i}}{k_{deg} + \mu} = \sum_j \frac{v_j}{k_{cat,\ i,\ j}} \tag{6}$$

Then, we set another two constraints for each complex, describing the degradation of proteins (degraded with rate $k_{deg}$) and dilution by growth (with rate $\mu$):

$$v_{cplx\ degradation,\ i} = v_{cplx\ formation,\ i} \times \frac{k_{deg}}{k_{deg} + \mu} \tag{7}$$

$$v_{cplx\ dilution,\ i} = v_{cplx\ formation,\ i} \times \frac{\mu}{k_{deg} + \mu} \tag{8}$$

#### 2.2 Protein translation

In the model, we describe protein translation as use of loaded amino acid-tRNAs ($tRNA^{AA}(AA)$) and retrieval of free tRNAs ($tRNA^{AA}$). First, free amino acids (AAs) are loaded onto respective tRNAs (Eq. 9), with a cost of 2 equivalents of ATP per amino acid. Then, loaded amino acid-tRNAs are used for protein translation according to the amino acid composition of the example protein *P00000* ($v_{syn,\ P00000}$) (Eq. 10), with $l_{P00000}$ being the peptide chain length. One cycle of peptide chain elongation costs 2 GTP, setting the total energetic cost of protein synthesis to be 4 NTPs per amino acid.

$$AA + tRNA^{AA} + ATP \rightarrow tRNA^{AA}(AA) + AMP + PP_i \tag{9}$$

$$\begin{aligned} v_{syn,\ P00000} : n_1\ tRNA^{AA_1}(AA_1) + \cdots + n_{20}\ tRNA^{AA_{20}}(AA_{20}) + 2l_{P00000}\ GTP + 2l_{P00000}\ H_2O \rightarrow \\ n_1\ tRNA^{AA_1} + \cdots + n_{20}\ tRNA^{AA_{20}} + 2l_{P00000}\ GDP + 2l_{P00000}\ P_i + 2l_{P00000}\ H^+ + P00000\_uf\_c \end{aligned} \tag{10}$$

##### Ribosome capacity coupling

We described the coupling between protein synthesis fluxes and ribosome formation as a function of peptide length. As we show in the Main Text, we estimated the ribosome parameters using quantitative proteomics data from [Kleijn et al., 2021]. The ribosomal peptide elongation rate was set to $k_{cat,\ ribo} = 10.5\ aa\ s^{-1} = 3.78 \times 10^4\ aa\ h^{-1}$, as also reported for *S. cerevisiae* by [Metzl-Raz et al., 2017]. The $k_{cat,\ ribo}$ is used to scale the ribosome demand to the protein synthesis fluxes. Also, [Metzl-Raz et al., 2017] have showed that the fraction of ribosomal proteins in the total proteome increases linearly with the growth rate $\mu$, with a offset value of $\phi_R^0$. We estimated the $\phi_R^0 \approx 0.0465\ g\ (g\ protein)^{-1}$ in *S. pombe* ($\phi_R^0 \approx 0.08$ for *S. cerevisiae*). As discussed in the Supplementary Note 6 of the *pcYeast7.6* model [Elsemman et al., 2021], in order to capture the offset in the model, we need to decompose this value into respective values for (a) truly inactive ribosomes $\phi_R^{0'}$ and (b) the ribosomes, required to synthesize the inactive ribosomes. Following the computation in [Elsemman et al., 2021], we computed the fraction $\phi_R^{0'} \approx 0.0306$. With the molecular weight of the protein component of the ribosome $MW_{P,\ ribo} = 1624.620\ g\ mmol^{-1}$, we compute the increment in the flux through ribosome complex formation reaction. Thus the general formulation of coupling constraint in Eq. 6 becomes:

$$\frac{k_{cat,\ ribo}[aa\ h^{-1}]}{k_{deg} + \mu} \times v_{cplx\ formation,\ ribosome} = \sum_{i \in cytoRibo} l_i[aa] \times v_{syn,\ i} + \frac{\phi_R^{0'} \times m_{prot}}{MW_{P,\ ribo}} \tag{11}$$

##### Translation factor demand

A number of translation initiation, elongation, and termination factors are needed for protein synthesis in ribosomes (Table 4). We thus considered different aspects of the translation for these groups of translation factors.

We first considered the translation initiation factors (eukaryotic initiation factors, eIFs). Their demand can be linked with the time, needed for ribosome to locate the initiation codon and fully assemble. For this, we first collected information on the length of the 5'-untranslated region (5'-UTR) from Pombase [Lock et al., 2019]. For mRNAs without a determined length of the 5'-UTR, we assumed the median value of 172 nucleotides. We assumed that ribosome scans the 5'-UTR region of the mRNA at the speed of $k_{cat,\ eIF} = 10\ nt\ s^{-1} = 3.6 \times 10^4\ nt\ h^{-1}$. Following that, we set the following coupling constraints for each of the individual complexes of translation initiation factors (Table 4):

$$\frac{k_{cat,\ eIF}[nt\ h^{-1}]}{k_{deg} + \mu} \times v_{cplx\ formation,\ eIF_i} = \sum_{j \in cytoRibo} l_{5'UTR,\ j}[nt] \times v_{syn,\ j} \tag{12}$$

Next, we coupled the demand for translation elongation factors (eEFs) in a similar manner. It should be noted that clear estimates of the turnover values of these factors are not available. Thus we reason that the $k_{cat,\ EF}$ value should be $k_{cat,\ eEF} \geq 10.5\ aa\ s^{-1}$. We can assume this because otherwise, the observed maximal speed of peptide elongation $k_{cat,\ ribo}$ would be considerably lower, as elongation is the longest phase in the translation process. Using this information, we set the coupling constraints for all individual complexes of translation elongation factors:

Table 4: Translation factors (and groups of), defined in the *pcPombe* model.

| Factor type | Cytosolic factors | Mitochondrial factors |
|---|---|---|
| Initiation factors | eIF2, eIF2B<br>eIF3<br>eIF4 | mIF |
| Elongation factors | eEF1<br>eEF2<br>eEF3 | mEF |
| Release factors | eRF | mRF |

$$\frac{k_{cat,\ eEF}[aa\ h^{-1}]}{k_{deg}+\mu} \times v_{cplx\ formation,\ eEF_i} = \sum_{j \in cytoRibo} l_j[aa] \times v_{syn,\ j} \tag{13}$$

Eukaryotic release factor (eRF) is the protein complex (eRF1 and eRF3), facilitating the dissociation of ribosomes (termination of translation) and release of fully translated peptide. The termination rate in *S. cerevisiae* is measured to be $k_{cat,\ eRF} = 0.15\ s^{-1} = 540\ h^{-1}$ [Shoemaker and Green, 2011].

$$\frac{k_{cat,\ eRF}[h^{-1}]}{k_{deg}+\mu} \times v_{cplx\ formation,\ eRF} = \sum_{i \in cytoRibo} v_{syn,\ i} \tag{14}$$

**Note on mitochondrial translation**

Mitochondria also have their own ribosomes, which translate a handful of protein species (including one subunit of the mitochondrial ribosome). We set coupling constraints, similar to the cytosolic ribosomes, to describe the demand of mitochondrial ribosomes:

$$\frac{k_{cat,\ mito\ ribo}[aa\ h^{-1}]}{k_{deg}+\mu} \times v_{cplx\ formation,\ mito\ ribosome} = \sum_{i \in mitoRibo} l_i[aa] \times v_{syn,\ i} \tag{15}$$

We chose to estimate the $k_{cat,\ mito\ ribo}$ value, rather than apply a "offset" value, as we did for cytosolic ribosomes. We thus used the estimate of mitochondrial ribosome elongation rate from *S. cerevisiae*, using quantitative proteomics data [Elsemman et al., 2021]. There, the value of $k_{cat,\ mito\ ribo} = 8\ aa\ s^{-1} = 2.8 \times 10^4\ aa\ h^{-1}$ has shown the best agreement to the experimental measurements. We also set up constraints to describe the use of mitochondrial translation factors, with similar expressions (Eqs. 12, 13, and 14) the same $k_{cat}$ values as for their cytosolic counterparts.

#### 2.3 Protein folding and transport

The unfolded protein species (synthesis described by Eq. 10) then have to be folded into "folded" (active) protein species. Optionally, these proteins first have to be transported to their target compartments. The folding reactions $v_{folding,\ P00000}$, catalyzed by chaperones (see coupling constraints, explained later), take the following form:

$$v_{folding,\ P00000} : P00000\_uf\_c \rightarrow P00000\_c \tag{16}$$

The transport reactions (transport demands explicitly modelled for mitochondria, through use of TIM/TOM translocase system) for the unfolded protein species **to** the target compartment (e.g. Golgi) and for the used protein species **out** of the compartment (except for mitochondria, see "Protein degradation") are as follows:

$$v_{import,\ P00000} : P00000\_uf\_c \rightarrow P00000\_uf\_g \tag{17}$$

$$v_{export,\ P00000} : P00000\_used\_g \rightarrow P00000\_used\_c \tag{18}$$

Table 5: Chaperones, defined in the *pcPombe* model.

| Chaperone | Coefficient | Target compartment(s) | $k_{cat}[h^{-1}]$ | Source |
|---|---|---|---|---|
| Hsp70<br>Hsp90 | 0.5<br>0.5 | Cytosol, Nucleus,<br>Vacuole | | [Lopez-Buesa et al., 1998]<br>(for *S. cerevisiae* Hsp70) |
| BiP | 1.0 | Golgi<br>ER | 78.0 | [Rosam et al., 2018] |
| mCp60 | 0.5 | Mitochondria | 57.0 | Same as for Hsp70 |

##### Chaperone coupling

We specified some chaperones in the model (Table 5) for protein folding. Some of these chaperones are specific for some compartments (such as BiP or mitochondrial Hsp class chaperon Mcp60), others were set to work in other compartments. Given that several chaperones could fold the same clients, we also attributed arbitrary coefficients $c_i$ ("workload sharing") to different chaperones. In the end, similarly to coupling constraints for ribosomes or translation factors, we described the coupling between folding reactions and chaperone use as follows:

$$\frac{k_{cat,\ chap_i}[h^{-1}]}{k_{deg}+\mu} \times v_{cplx\ formation,\ chap_i} = c_i \sum_{j \in chap_i} v_{folding,\ j} \tag{19}$$

##### TIM/TOM translocase coupling

We modeled protein transport to mitochondria explicitly due to the fact that mitochondria are metabolically active organelles and also they impose proteome limits on cellular metabolism.
Translocation of proteins to mitochondria begins by pulling the unfolded peptide through the channels, formed by the outer- and inner membrane translocases (denoted as $t_i$ in equations). Three kinds of translocases are defined in the model: TOM, the outer membrane translocase, TIM22, and TIM23 (both inner membrane translocases). In the *pcYeast7.6* model [Elsemman et al., 2021], we estimated the turnover value of $k_{cat,\ t} = 3\ min^{-1} = 180\ h^{-1}$, and set the following coupling constraint for each translocase complex:

$$\frac{k_{cat,\ t}[h^{-1}]}{k_{deg}+\mu} \times v_{cplx\ formation,\ t_i} = \sum_{j \in mito} v_{import,\ j} \tag{20}$$

#### 2.4 Protein degradation

Used protein species are degraded to individual amino acids either in cytosol (cytosolic proteins and proteins from compartments other than mitochondria) or in mitochondria. In the first case, degradation is catalyzed by the proteasome complex, in the latter - by mitochondrial protease *PIM1*. In both cases, proteins are hydrolyzed to individual amino acids, consuming an estimated 1.35 ATP per amino acid released (estimate for *S. cerevisiae* from [Hong et al., 2012]).

$$\begin{aligned} v_{degradation,\ P00000} : P00000\_used\_c + 1.35n\ ATP + 2.35n\ H_2O \rightarrow \\ n_1\ AA_1 + \cdots + n_{20}\ AA_{20} + 1.35n\ ADP + 1.35n\ P_i + 1.35n\ H^+ \end{aligned} \tag{21}$$

##### Proteasome and *PIM1* coupling

As mentioned in the section above, all but mitochondria-targeted proteins are exported as their used forms from their target compartments and degraded in the cytosol (Eq. 18). In the model, we couple cytosolic protein degradation with the use of proteasomes. A proteasome is a large protein complex which unfolds and degrades proteins into short peptides and/or free amino acids. Similarly, in mitochondria, both unfolding and degradation happen, the latter catalyzed by PIM1, which is the homolog of prokaryotic *Lon* protease. We considered the turnover values of both proteasome $k_{cat,\ proteasome} = 2.3\ min^{-1} = 138\ h^{-1}$ [Peth et al., 2013] and PIM1 $k_{cat,\ PIM1} = 0.26\ s^{-1} = 936\ h^{-1}$ [Patterson-Ward et al., 2007] from literature, and set the coupling constraints in both cases:

$$\frac{k_{cat,\ proteasome}[h^{-1}]}{k_{deg}+\mu} \times v_{cplx\ formation,\ proteasome} = \sum_{j \in proteasome} v_{degradation,\ j} \quad (22)$$

and

$$\frac{k_{cat,\ PIM1}[h^{-1}]}{k_{deg}+\mu} \times v_{cplx\ formation,\ PIM1} = \sum_{j \in PIM1} v_{degradation,\ j} \quad (23)$$

### 3 Compartment-specific proteome constraints

In the model, we use three sets of different compartment-specific proteome constraints (also called "capacity constraints"), by the parameter of the proteins which is used to formulate the constraint: protein mass (molecular weight), protein crossection surface area, and protein volume. In the following, we describe these constraints in more detail.

#### 3.1 Proteome mass constraints

##### 3.1.1 Total proteome mass constraint

The proteome mass constraint is an equality constraint, i.e. its expression must always satisfy the right-hand-side (RHS). This constraint therefore sets the protein density in the dry biomass to be consistent with experimental measurements. For the *S. pombe* wild-type cells, [de Jong-Gubbels et al., 1996] reported that the bulk protein mass fraction the dry weight $f_{p,biomass}(\mu)$ is relatively independent of growth rate $\mu$ ($f_{p,biomass} = 0.43\ (g\ protein)\ gDW^{-1}$). Following these measurements, we set the total proteome mass to equal:

$$\sum_i \frac{v_{syn,i}}{k_{deg}+\mu} \times MW_i\ [\frac{g\ protein}{mol}] \times 10^{-3}\ [\frac{mol}{mmol}] = f_{p,biomass}\ [(g\ protein)\ gDW^{-1}] \quad (24)$$

##### 3.1.2 Mitochondrial maintenance coupling

Previously we described the mitochondria-specific maintenance reaction (Section 1.3), which we used to impose extra costs of maintaining mitochondria, which are not explicitly described in the model. The energetic costs in the maintenance reaction account for the ATP hydrolysis, used in, among others, translocation of proteins into mitochondria (Section 2.3). We specified the costs of mitochondrial maintenance per gram mitochondrial protein, by coupling the flux through the maintenance reaction to the mass of mitochondrial proteins:

$$\sum_{i \in mito} \frac{v_{syn,i}}{k_{deg}+\mu} \times MW_i\ [\frac{g\ protein}{mol}] \times 10^{-3}\ [\frac{mol}{mmol}] = v_{Mitochondrial\ maintenance} \quad (25)$$

##### 3.1.3 Unspecified protein constraint

In the model, not all proteins, present in the proteome of *S. pombe* are defined. These proteins can be without enzymatic annotation (e.g. structural proteins), not directly metabolic (e.g. signaling proteins) or simply be not annotated. However, our previously published proteomics data ([Elsemman et al., 2021]) suggests that more than 2-2.5 thousand protein species can be robustly quantified using label-free mass spectrometry-based proteomics, which is a substantially higher number than the number of protein species, predicted to be expressed by the model. Moreover, at low growth rates, many proteins are expressed in excess, and we always compute the minimal requirement of these (see 2.1). In order to account for the protein expression, which are not explicitly defined in the model, we created an artificial protein species, *PDUMMY*, which we call the "unspecified protein" (UP). Based on the proteomics measurements in glucose batch conditions [Kleijn et al., 2021], proteins that are not represented in the model, occupy ca. 25% of the total protein mass. Thus we set the minimal level of expression of the UP at every growth rate $\mu$, which corresponds to the proteome mass fraction $\phi_{UP}$. As described previously (Section 1.3), we specified some maintenance requirements for the mitochondria. Following suit, the unspecified protein demand thus was also split into two constraints, one for the cytosolic ("general") UP expression demand:

$$\frac{v_{folding,\ cyto,\ UP}}{k_{deg}+\mu} \times MW_{UP}\ [\frac{g\ UP}{mol}] \times 10^{-3}\ [\frac{mol}{mmol}] \geq \phi_{UP}\ [(g\ UP)\ (g\ protein)^{-1}] \times f_{p,biomass}(\mu) \quad (26)$$

Alternatively, can we formulate the (Eq. 26) so that we do not have to explicitly input the value of $f_{p,biomass}(\mu)$ into the constraint expression:

$$\sum_{i \in proteins} \frac{v_{syn,\ i}}{k_{deg}+\mu} \times MW_i\ [\frac{g\ protein}{mol}] - \frac{1}{1-\phi_{UP}} \times MW_{UP}\ [\frac{g\ UP}{mol}] \times v_{folding,\ cyto,\ UP} \leq 0 \quad (27)$$

Then, the mitochondrial UP experession demand (determined to be 32% of total mitochondrial proteome) is set as follows:

$$\sum_{i \in mito\ proteins} \frac{v_{folding,\ mito,\ i}}{k_{deg}+\mu} \times MW_i\ [\frac{g\ protein}{mol}] - \frac{1}{1-\phi_{UP,\ mito}} \times MW_{UP}\ [\frac{g\ UP}{mol}] \times v_{folding,\ mito,\ UP} \leq 0 \quad (28)$$

#### 3.2 Protein area and volume constraints

##### Constants and bionumbers for conversion

These constraints are expressed by area/volume per cell, therefore, some conversion units are needed for establishing relations between parameters of proteins and cells.
First, we compute the volume of a protein molecule, based on its molecular weight. Following [Erickson, 2009], we assume the protein to be globular (to form a sphere when folded). Then, we compute the minimal radius of the protein molecule with that molecular weight ($r_{min}$), using the following empirical relation:

$$r_{min}\ [nm] = 0.066 \times MW^{\frac{1}{3}} \quad (29)$$

Further, assuming the sphere-shaped protein molecule, we can compute its crossection area (area of a disc with the radius $r_{min}$) and volume (also using the radius $r_{min}$):

$$A_{peptide}\ [nm^2] = \pi \times r_{min}^2 \quad (30)$$

$$V_{peptide}\ [nm^3] = \frac{4}{3} \times \pi \times r_{min}^3 \quad (31)$$

Next, we provide the relations needed to compute the parameters (volume, surface area etc.) of the cell. Since all fluxes in the model are scaled to gram dry cell weight ($gDW$), it is handy to convert the units of cell volume to cell dry biomass. [Odermatt et al., 2021] have reported that 1 gDW of cells occupy a volume of roughly 3.55 mL, and this number is relatively stable at different growth rates:

$$V_{gDW} = 3.55\ mL\ gDW^{-1} \quad (32)$$

Even though the volume-dry weight relationship is fixed, it is known that faster-growing *S. pombe* cells are larger. We used experimental data from [Vraná, 1983, Sveiczer et al., 1996, Hayles and Nurse, 2001] to compute growth rate $\mu$-dependent cell volume:

$$V_{cell}\ [\mu m^3] = 152.318 \times \mu + 35.93 \quad (33)$$

Assuming a cylinder-like cell (resembling the natural shape of *S. pombe*) with a radius of $r_{cell} = 1.75\ \mu m$ [Hayles and Nurse, 2001], we can compute the length of such a cell, and, subsequently, its surface area.

$$A_{crosssection} = r_{cell}^2 \times \pi \quad (34)$$

$$l_{cell}\ [\mu m] = \frac{V_{cell}}{A_{crosssection}} \quad (35)$$

$$A_{cell}\ [\mu m^2] = (2\pi \times r_{cell}) \times l_{cell} \quad (36)$$

Following Eqs. 32 and 33, a number of cells in a gram dry weight can be computed:

$$N_{cells} = \frac{V_{gDW} \times 10^{-6}\ [\frac{\mu m^3}{mL}]}{V_{cell} \times 10^{-18}\ [\frac{m^3}{\mu m^3}]} \quad (37)$$

##### 3.2.1 Plasma membrane area constraints

We use the following constraint expression to define the protein capacity in the plasma membrane of the cell. In the model, we assume that 30% of the plasma membrane ($PM$) area is accessible for proteins to occupy ($f_{PM}$). Then, for two specific groups of proteins, transporters of carbon sources ($PMC$) and nitrogen sources ($PMN$), we formulate two separate constraints, assuming these transporters can occupy 5.0% ($f_{PMC}$) and 3.0% ($f_{PMN}$)of plasma membrane surface area, respectively.
First we compute the sum of the molecular weight of the protein complex, formed in the plasma membrane, based on the molecular weight ($MW$) and stoichiometry ($n$) of the proteins, forming the complex:

$$MW_{cplx} = \sum_{i \in cplx} n_i \times MW_i \quad (38)$$

Using Eqs. 29 and 30, the crossection area of this protein complex (per $mmol$ of complex) is computed as follows:

$$A_{mmol\ cplx} = A_{peptide}(MW_{cplx})\ [nm^2] \times 10^{-6}\ [\frac{\mu m^2}{nm^2}] \times 6.022 \times 10^{20}\ [mmol^{-1}] \quad (39)$$

Next, we can compute the surface area the $mmol$ of this protein complex would occupy per single cell (based on the cell number computation from Eq. 37):

$$A_{mmol,sc} = \frac{A_{mmol\ cplx}}{N_{cells}} \quad (40)$$

Finally, we formulate the constraint of available plasma membrane area per single cell:

$$\sum_{i \in compartment} \frac{v_{cplx\ formation,\ i}}{k_{deg} + \mu} \times A_{mmol,sc} \leq f_{compartment} \times A_{cell}\ [\mu m^2\ cell^{-1}] \quad (41)$$

Note that here we sum not over protein translation fluxes ($v_{syn}$), but through complex formation fluxes ($v_{cplx\ formation,\ i}$). This is a precaution to make sure that only these proteins, which form complexes, attributed to plasma membrane (the target compartment remains cytosol), are counted. In the mitochondrial volume constraint (see below), sum is also computed not through $v_{syn}$ reactions due to the same issue.

##### 3.2.2 Mitochondrial capacity constraint

Similarly to the previously described constraint, we compute the volume of a single peptide and scale it per $mmol$ and per single cell. Since there is no ambiguity of whether the protein should be attributed to mitochondrial capacity or not (see the Eq. 44), we sum over individual proteins rather than their complexes.

$$V_{mmol} = V_{peptide}\ [nm^3] \times 10^{-9}\ [\frac{\mu m^3}{nm^3}] \times 6.022 \times 10^{20}\ [mmol^{-1}] \quad (42)$$

$$V_{mmol,sc} = \frac{V_{mmol}}{N_{cells}} \quad (43)$$

We formulate a mitochondrial capacity constraint in the following way, summing the fluxes through protein folding reactions in mitochondria ($v_{folding,\ mito,\ i}$), with the estimated volume, allocated for mitochondrial proteins $V_{mito\ proteins} = 0.8\ \mu m^3\ cell^{-1}$:

$$\sum_{i \in Mito} \frac{v_{folding,\ mito,\ i}}{k_{deg} + \mu} \times V_{mmol,sc} \leq V_{mito\ proteins}\ [\mu m^3\ cell^{-1}] \quad (44)$$
